# The Use of Distributions in SBML Models

**DOI:** 10.1101/015503

**Authors:** Lucian Smith, Herbert M. Sauro

## Abstract

In this technical note we describe modifications to Antimony (Biochemical model specification language) that allows modelers to use the SBML distributions package. In addition the article describes best practice for using distributions, including when they should and should not be used.

## INTRODUCTION

With the SBML L3 Package “Distributions”, [1] it has become possible to exchange SBML [2] models with embedded distributions. This allows modelers to encode stochastic aspects of the system they are modeling directly in the model.

There are two ways to use the Distributions package. The first uses new constructs that allow one to, essentially, annotate any mathematical element in an SBML file with information about a distribution associated with that element. This allows new types of analysis of a model without changing the original model at all. For example, one could annotate all input parameters with standard deviations, and do an error analysis of simulator output. Or one could annotate parameters as coming from particular distributions, then run a series of simulations, each drawing a new value for those parameters from those distributions.

The second method extends the SBML ‘function definition’ construct. An SBML function definition takes arguments and returns the result of a computation on those inputs. The Distributions package adds the ability to define a new function definition that again takes arguments, but which returns a draw from a distribution defined by those inputs. Because this use defines a draw from a distribution, it should not generally be used in continuous contexts. In SBML, that means not using them:

- In rate rules or kinetic laws: These constructs define how variables change in time. Continually drawing from a distribution would mean that over any given time interval, all possible draws from that distribution would be sampled, including (in many cases) positive and negative infinity. This situation is obviously intractable.
- In assignment rules and algebraic rules: These constructs define the value of SBML variables over all times. Again, since all possible draws from that distribution are possible over any given time interval, this would make normal SBML model analysis impossible.
- In event triggers: This is a particularly important situation to avoid. When an event trigger changes in value from *false* to *true*, the event is fired, causing its assignments to be carried out. If all possible draws from a distribution are possible over any given time interval, this would mean that a typical trigger would literally fire an infinite number of times for every time step. At the very least, it would mean that any time it was remotely possible that a trigger would fire, it would do so.

Note that with additional information such as autocorrelation values or full conditional probabilities, the above situations could become tractable once more, as the distribution would be defined over time as a random walk, not as pure white noise.

All other uses of MathML in SBML are viable places to use the new distribution function constructs:

- In initial assignments: An initial assignment is calculated exactly once at the beginning of a simulation. A model with a draw from a distribution in an initial assignment, then, would behave differently from one run to another, but the call would not exert any other stochastic influence on the simulation once the run had started.
- In ‘event’ assignments, delays, and priorities: As mentioned above, an event in SBML is triggered when the evaluation of its trigger function changes from *false* to *true*. After an optional delay, any assignments it might have are executed. If multiple events are to be executed at the same moment in time, the order in which they do so is determined by their relative priorities, with events with higher priorities being executed first. Because the calculation of the delay, calculation of relative priorities, and calculation of the value to be assigned all take place at discrete moments in the simulation, all three are viable places for calls for draws from distributions. These models will behave stochastically regardless of their initial state, with a single event potentially behaving differently if called multiple times within a single simulation.

The behavior of a draw from a distribution in an initial assignment is straightforward to understand, but the complexity of SBML events allows for a variety of opportunities for using distributions that are worth exploring in greater detail. The use of distributions in delays

To begin with, modeling a situation that might at first seem solvable by putting a draw from a distribution in a Trigger (which we have seen is actually intractable) might instead be better solved by putting a draw from a distribution in a Delay. Suppose we are modeling a neural depolarization event. Once certain conditions are met, there is a certain percentage chance of the event firing per second. This can be modeled by calculating the distribution of times that it is likely to take until the event takes place: an exponential distribution is likely to be a good candidate for this situation. Giving the event a Trigger of the conditions to be met, and a Delay of a draw from that distribution will exactly model the random depolarization of the neuron.

Of course, in many situations, it is not the case that the neuron, once certain conditions are met, is guaranteed to depolarize at some point in the future. Instead, those conditions must continually be true until the depolarization event happens. This is the situation for which the *persistent* attribute on the Trigger was created. If this attribute is set to *false*, the event, once triggered, must fire; it is only a matter of when this will happen. If it is set to ‘true’, however, the trigger conditions must not only change from *false* to *true*, but those conditions must not change back to *false* before the event fires. Should the conditions change, the event will no longer fire, regardless of the time it was originally scheduled to do so.

To put this in concrete terms, let us imagine a model of a particular neuron. If the concentration of S1 goes above 5, a depolarization event is likely to happen, taking an average of 2 seconds to do so, which will reset the concentration of S1 to zero. However, if the concentration of S1 goes below 5, depolarization can no longer take place. This can be modeled by the following SBML event, written here in Antimony shorthand:

~~~
depolarization: at exponential(0.5) after S1 > 5, persistent=true: S1=0
~~~

Or, in SBML XML:

~~~
<event id="depolarization" useValuesFromTriggerTime="true">
  <trigger initialValue="true" persistent="true">
    <math xmlns="http://www.w3.org/1998/Math/MathML">
      <apply>
        <gt/>
        <ci> S1 </ci>
        <cn type="integer"> 5 </cn>
      </apply>
    </math>
  </trigger>
  <delay>
    <math xmlns="http://www.w3.org/1998/Math/MathML">
      <apply>
        <ci> exponential </ci>
        <cn> 0.5 </cn>
      </apply>
    </math>
  </delay>
  <listOfEventAssignments>
    <eventAssignment variable="S1">
      <math xmlns="http://www.w3.org/1998/Math/MathML">
        <cn type="integer"> 0 </cn>
      </math>
    </eventAssignment>
  </listOfEventAssignments>
</event>
~~~

(The call to the *exponential* function here will be to a ‘Distributions’-extended function definition that takes the rate parameter as its argument.)

The reason this setup works is because the exponential function is ‘memoryless’, that is, if the event has not fired at a certain point after being triggered, the probability of it firing in the next n seconds is the same as if the event had just been triggered. Thus, if S1 was greater than 5 for three seconds and then went below five again, if the chosen delay was 4.2, it is sufficient that 4.2 > 3 and say that the depolarization event did not happen during that three-second window. It is not necessary, should S1 again go above 5, to remember that 1.2 seconds were ‘left over’ from the previous delay: the exact same distribution will be obtained if a new draw from the exponential for the delay is chosen the second time the trigger condition is met.

## THE USE OF DISTRIBUTIONS IN PRIORITIES

A Priority in an SBML Event allows a modeler to define, when two events are to be executed at the same moment in time, which event execution should take place first. If one event has a higher priority than the other, it is executed first, and if two events have the same priority, they are to be executed in a random order with respect to one another.

The use of distributions in Priority math allow the modeler to model situations where a single situation has two potential outcomes, with one more likely to happen than the other. To create a very simple case, where when S1 gets above a certain threshold, it is either converted to S4 (25% probability) or S5 (75% probability):

~~~
S1toS4: at S1 > 5, persistent=true, priority = uniform(0,1): S4 = S1, S1 = 0;
S1toS5: at S1 > 5, persistent=true, priority = uniform(0,2): S5 = S1, S1 = 0;
~~~

Here, both events are triggered by the same condition, making them simultaneous. 50% of the time, the draw from the uniform distribution of second event will be greater than 1, necessarily giving it the higher priority. The remaining 50% of the time, both priorities will be chosen from the same range of values, giving both events a 25% chance of being higher than the other.

Note that we must set the ‘persistent’ attribute to *true*, so that when either event is executed and the level of S1 returns to zero, the trigger condition for the other event is no longer *true*, meaning it is not to be executed. These conditional probabilities can be calculated from model parameters, as well. For this next model, we imagine a cell with potassium and sodium channels that each might open when the total ion concentration in the cell gets too high, with each more likely to be the one to open if the concentration of its corresponding ion is higher than the other:

~~~
// The channels themselves:
Kchannel:      K -> K_ext; Kon*K;
Nachannel: Na -> Na_ext; NaOn*Na;
~~~

~~~
// Turning one of the channels on:
at (K+Na) > 8 && Kon == 0, persistent = true, priority=uniform(0,Na): NaOn = 1;
at (K+Na) > 8 && NaOn == 0, persistent = true, priority=uniform(0,K): KOn = 1;
~~~

~~~
// Turning the channels back off:
at 5 after Kon > 0:      Kon=0;
at 5 after NaOn > 0: NaOn=0;
~~~

~~~
// Initial conditions:
Kon = 0;
NaOn = 0;
~~~

## THE USE OF THE SAME DISTRIBUTION IN MULTIPLE CALCULATIONS

When the same draw from a distribution is desired in multiple contexts, it may be necessary to set up multiple events: one to assign the value from the distribution to a variable, and one or more called directly afterwards that use it. For an example, let us take a model of cell division that is using amounts (instead of concentrations) of species. If the cell were to always divide exactly in half, one could use an event like the following (again in Antimony shorthand):

~~~
cellDivision: at (telo == 2): S1 = 0.5*S1, S2 = 0.5*S2, S3 = 0.5*S3;
~~~

However, if one wanted to have the division percentage be a normal distribution, one could not simply do:

~~~
cellDivision: at (telo == 2): S1 = normal(0.5,0.01)*S1,
S2 = normal(0.5,0.01)*S2,
S3 = normal(0.5,0.01)*S3;
~~~

Because each call to “normal(0.5,0.01)” would result in a separate draw from the distribution, de-synchronizing the amounts of S1, S2, and S3. Instead, one would need to set up a first event that performed a draw from a distribution and stored it, and then a second event that used that value. Here, we give two events the same trigger, meaning that they will always be simultaneous, and then give the first a higher priority than the second, meaning that its event assignments will be carried out before the other event assignments. However, to let that new value be used in the second event, the second event needs to be set to not use the values in its assignments from the time it was triggered, but from the time the assignment is to be made. In Antimony, this is set by using the `fromTrigger` keyword, which is translated to SBML as the event attribute “useValuesFromTriggerTime”:

~~~
cellDivisionA: at (telo == 2), priority = 10: divisionPercent = normal(0.5, 0.01);
cellDivisionB: at (telo == 2), priority = 1, fromTrigger = false:
S1 = divisionPercent*S1,
S2 = divisionPercent*S2,
S3 = divisionPercent*S3;
~~~

Alternatively, one could make a slightly simpler model that would have the same effect by setting ‘divisionPercent’ with an initial assignment to be used the first time the event is fired, and have another event assignment that re-set ‘divisionPercent’ so it would have a new value the next time it was fired:

~~~
divisionPercent = normal(0.5, 0.01);
cellDivision: at (telo == 2): S1 = divisionPercent*S1,
S2 = divisionPercent*S2,
S3 = divisionPercent*S3; divisionPercent = normal(0.5, 0.01);
divisionPercent = normal(0.5, 0.01);
~~~

This would not be possible, however, in a situation where the division percentage was dependent on the conditions of the model at the time of division. In that case, the priority approach would have to be taken. Truncated distributions

In some cases, a given distribution may reasonably approximate the true distribution being modeled, but may include values that do not make sense for the system. In the above case, though a normal distribution centered at 0.5 may be reasonable approximation of the vagaries of cell division around the mean, it would be possible to draw values of less than zero or greater than one, which do not make physical sense for this process. While some other distribution might be found with natural boundaries that match the physical limits, it is sometimes simpler to instead define the distribution as being truncated. Antimony provides a shorthand way of defining these truncations with a different form of the function. If we believe that no modeled cell division could physically take place at any ratio lower than 9:1, the last model above, then, would become:

~~~
divisionPercent = truncatedNormal(0.5, 0.01, 0.1, 0.9);
cellDivision: at (telo == 2): S1 = divisionPercent*S1,
S2 = divisionPercent*S2,
S3 = divisionPercent*S3;
divisionPercent = truncatedNormal(0.5, 0.01, 0.1, 0.9);
~~~

The relevant change corresponding SBML model is the addition of ‘truncationLowerInclusiveBound’ and ‘truncationUpperInclusiveBound’ elements in the UncertML, as well as new arguments to set those values.:

~~~
<distrib:drawFromDistribution>
  <distrib:listOfDistribInputs>
    <distrib:distribInput distrib:id="mean" distrib:index="0"/>
    <distrib:distribInput distrib:id="stddev" distrib:index="1"/>
    <distrib:distribInput distrib:id="lowlimit" distrib:index="2"/>
    <distrib:distribInput distrib:id="uplimit" distrib:index="3"/>
  </distrib:listOfDistribInputs>
  <UncertML xmlns="http://www.uncertml.org/3.0">
    <NormalDistribution definition="http://www.uncertml.org/distributions">
      <mean>
        <var varId="mean"/>
      </mean>
      <stddev>
        <var varId="stddev"/>
      </stddev>
      <truncationLowerInclusiveBound>
        <var varId="lowlimit"/>
      </truncationLowerInclusiveBound>
      <truncationUpperInclusiveBound>
        <var varId="uplimit"/>
      </truncationUpperInclusiveBound>
    </NormalDistribution>
  </UncertML>
</distrib:drawFromDistribution>
~~~

Note that a truncated distribution does not mean that the probability density that would have fallen outside of the truncation is assigned to the borders of the truncation. Instead, the probability density of the entire distribution must be rescaled so that the total area is once again 1.0. This means that if one drew a value of ‘-0.3’ from the non-truncated version of the above distribution, one could not simply return 0.1 (the lower limit) and claim that this constituted a draw from the truncated distribution. In that situation, an entirely new draw from the distribution must be made, or a method of drawing directly from the truncated distribution itself must be devised.

## SUMMARY OF SUPPORTED DISTRIBUTIONS

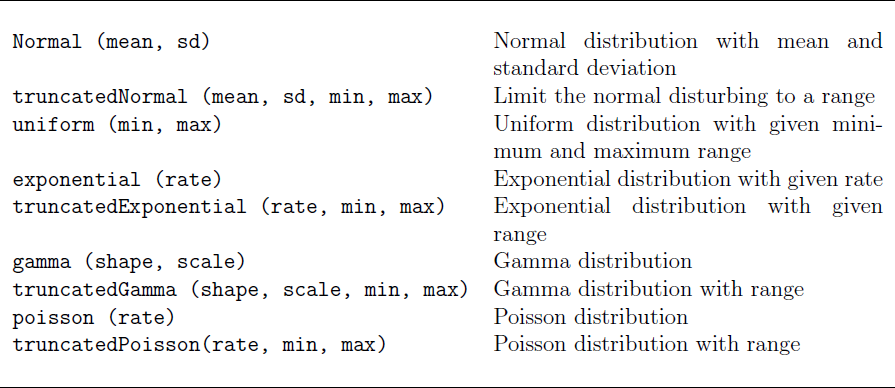

## ACKNOWLEDGMENTS

Herbert M. Sauro acknowledges support from NIH grant R01 GM081070. The content is solely the responsibility of the authors and does not necessarily represent the official views of the National Institutes of Health.

